# A conserved upstream element in the mouse *Csf1r* locus contributes to transcription in hematopoietic and trophoblast cells

**DOI:** 10.1101/2025.08.05.668600

**Authors:** Emma Maxwell, Isis Taylor, Cameron Flegg, Yajun Liu, Ginell Ranpura, Fathima Nooru Mohamed, Emma K Green, Stephen Huang, Jintao Guo, Ngari Teakle, Kamil A. Sokolowski, Katharine M. Irvine, David A. Hume, Sebastien Jacquelin

## Abstract

Expression of the *Csf1r* gene in mice is restricted to cells of the mononuclear phagocyte system and placental trophoblasts. A conserved element (*Csf1r* upstream regulatory element A, CUREA) in the mouse *Csf1r* locus contains transcription start sites utilised by trophoblasts and osteoclasts and an enhancer essential for expression of multicopy transgenic reporters in most tissue macrophages. Here we describe the impact of deletion of CUREA in the mouse genome, on the background of a knock-in *Csf1r*-FusionRed reporter. By contrast to the essential function in transgene expression, CUREA deletion had no effect on expression of FusionRed or differentiation of blood monocytes or tissue resident macrophages. The deletion reduced *Csf1r* mRNA in hematopoietic stem cells and committed myeloid progenitors (MPP3) leading to a subtle differentiation delay and also had a significant impact on microglial phenotype in the brain and the differentiation of osteoclasts. The expression of FusionRed in placenta confirmed expression of CSF1R in trophoblasts. 5’RACE analysis demonstrated that the effect of CUREA deletion on *Csf1r* transcription in placenta was overcome by the use of cryptic upstream transcription start sites. We conclude that CUREA is a regulatory element controlling *Csf1r* transcription. The function overlaps with other enhancers identified in the locus and is therefore partly redundant.

**Key Points:**

- A regulatory element (CUREA) in the mouse *Csf1r* locus has both promoter and enhancer activity.
- Germ-line deletion of CUREA impacts differentiation of marrow progenitors, microglia, osteoclasts and placental trophoblasts.

## Introduction

Differentiation, proliferation and survival of cells of the mononuclear phagocyte system (MPS) depends upon signaling from the macrophage colony stimulating factor receptor (CSF1R) initiated by the two ligands, CSF1 and interleukin 34 (IL34). Homozygous recessive mutations of *CSF1R* in mouse, rat and human lead to the loss of most tissue resident macrophage populations including bone resorbing osteoclasts and pleiotropic effects on postnatal development ^1^. CSF1R is expressed in the earliest detectable macrophages in the yolk sac in vertebrates and is restricted to the MPS lineage throughout embryonic and adult development ^2–4^. In mammalian extraembryonic tissues, *Csf1r* mRNA is also expressed in the ectoplacental cone and in placenta ^5,6^ transcribed from a separate trophoblast-specific promoter ^7,8^. Our group has studied transcriptional regulation of the mouse *Csf1r* locus as a model to understand MPS lineage commitment and differentiation (reviewed in ^9^). The major myeloid *Csf1r* promoter is purine-rich and lacks a TATA box. Transcription initiation occurs at a cluster of sites and depends upon binding of the macrophage-specific transcription factor, PU.1 (SPI1). We characterised a highly conserved intronic enhancer (the Fms intronic regulatory element, FIRE) that was absolutely required for expression of a multicopy reporter transgene ^8^. Deletion of the FIRE sequence in the mouse genome (*Csf1r*^ΔFIRE/ΔFIRE^) led to selective loss of macrophages in the brain (microglia) and some other tissues but in other locations, *Csf1r* mRNA was expressed and tissue macrophages were unaffected ^10^. Chromatin analysis revealed multiple additional candidate enhancers aside from FIRE that may contribute to the apparent redundancy ^10^.

We focused on an additional conserved regulatory element immediately upstream of the major macrophage transcription start sites (TSS) and proximal promoter in mice that we have called the CSF1R-upstream regulatory element A (CUREA). This element includes the transcription initiation sites active in mouse trophoblasts ^8^. During macrophage lineage differentiation from bone marrow progenitors, the 500bp promoter region including CUREA is in open chromatin prior to the intronic enhancer ^11^. In human hematopoietic progenitors CUREA can be classify as active enhancer based upon epigenetic histone modifications, H3K4me1 and H3K27ac ^12^. In multiple ATAC-seq data sets derived from mouse progenitors and isolated tissue macrophages, CUREA is located within a distinct peak of open chromatin ^13–16^. The function of CUREA was analysed by the generation of transgenic reporter mice using a promoter that included FIRE but lacked CUREA. In a series of *Csf1r-*EGFP transgenic lines, CUREA deletion abolished expression in placenta and in most tissue-resident macrophages ^17^. In the MacBlue line (*Csf1r*-Gal4VP16/UAS-ECFP) the *Csf1r* promoter with CUREA deleted directs expression of a transcription factor that in turn drives expression of cyan fluorescent protein (ECFP). The ECFP reporter is not expressed in placenta but is highly expressed in embryo macrophages and retained in the majority of microglia ^18,19^. It is also detected in blood monocytes but either absent or rapidly down-regulated in most tissue-resident macrophages ^19–21^. Here we have deleted CUREA in the mouse germ line and present an analysis of the selective impact on *Csf1r* expression and differentiation in macrophage and trophoblast lineages. The results indicate that CUREA contributes to transcriptional regulation in the macrophage lineage but is not essential for tissue-resident macrophage or trophoblast differentiation.

## Methods

### Generation of *Csf1r T2A-FRed* ^ΔCUREA/ΔCUREA^ mice and animal breeding

A ribonucleoprotein (RNP) complex comprised of EnGenSpy-Cas9 (NEB cat: M0646T) NLS and two synthetic SgRNAs (sequence provided in **Supp table 1)** targeting CUREA was microinjected into C57BL/6J/ *Csf1r T2A-FRed^+/+^* ^4^ zygote. Correct targeting was confirmed by PCR amplification and sequencing. For routine genotyping, the ΔCUREA allele was detected by PCR using primers 5’ TGCTATAGGAGGCTCATTACTGT 3’ and 5’ TCTTCCTCTGGTGTGTGGTG3’ and GoTaq Green (Promega). PCR conditions were 98°C 120 seconds, followed by 30 cycles at 98°C 30 seconds, 55°C for 30 seconds, 72°C for 30 seconds, then 72°C for 120 seconds. The *Csf1r*-Fred allele was genotyped as previously described ^4^. Mice were bred and maintained in a specific pathogen free facility within the Translational Research Institute. All studies were approved by the Animal Ethics Committee of the University of Queensland. Mice were housed in individually ventilated cages with a 12 h light/dark cycle, and food and water available *ad libitum*.

### Promoter capture

5′ RACE was performed using an NEB kit (NEB #M0466). To increase sensitivity and specificity Oligo dT was used as the primer for first strand synthesis followed by a nested PCR with gene-specific primer. cDNA was diluted 20x then PCR amplified using Q5 Hot Start High-Fidelity Master Mix (2X) (NEB #M0494) and primers; 5’ CATTGCAAGCAGTGGTATCAAC 3’ (template switch oligonucleotide, TSO) and 5’CTGACCATGCCAAACTGTGGCCAGCAGC 3’ (*Csf1r*-specific). PCR products were purified using a Min-Elute PCR Purification kit (Qiagen). 1ul of template was used for the second PCR using a nested *Csf1r* primer; 5’ATCGCAGGGTCACCGTTTCACCCGGC 3’ in combination with either TSO or primers F1.1 and F1.2 corresponding to previously identified 5’ ends of trophoblast and osteoclast specific *Csf1r* transcripts respectively ^17^. The PCR cycle was 98°C 120 seconds, followed by 30 cycles at 98°C 30 seconds, 58°C for 30 seconds, and 72°C for 30 seconds, then 72°C for 5 minutes.

mRNA quantitation by qRT-PCR, histochemistry, immunohistochemistry, tissue disaggregation, immunofluorescence staining, flow cytometry, cell isolation and culture, microCT and image analysis were performed using standard methods. Details and sources of reagents are provided in Supplementary Text.

## Results

### Generation of *Csf1r T2A-FRed* ^ΔCUREA/ΔCUREA^ Mice

To enable an analysis of the impact of CUREA deletion on the CSF1R expression and the distribution of CSF1R^+^ macrophages in tissues, the deletion was made in homozygous *Csf1r*-FusionRed mice which have a knock-in T2A-FusionRed reporter ^4^. A ribonucleoprotein (RNP) complex comprising EnGenSpy-Cas9 NLS and two synthetic SgRNAs targeting CUREA was microinjected in C57BL/6J/ *Csf1r T2A-FRed^+/+^* zygote. SgRNAs targeting CUREA were designed and selected using the Sanger WTSI Web site (http://www.sanger.ac.uk/htgt/wge/). Successful deletion was confirmed as shown in **Supp-Figure1A**. Genotyping was performed with primers flanking the deleted region (**Supp-Figure1B, C**). Mice with the desired deletion were crossed with wild type C57BL/6J. The progeny were interbred to generate homozygous mutants (CUREA^-/-^). CUREA^-/-^ mice were weaned at expected Mendelian frequency, were healthy, fertile and did not present any of the deleterious phenotypes observed in CSF1R^-/-^ or CSF1R^op/op^ mice ^22^.

### CUREA regulates *Csf1r* expression in hematopoietic stem and progenitor cells

We first investigated the distribution of hematopoietic stem cells and progenitors (HSPCs) in the bone marrow of CUREA^-/-^ and wild type (WT) mice (**Figure 1A**). There was a relative accumulation of LT-HSCs and ST-HSCs in CUREA^-/-^ bone marrow and a corresponding decrease in multipotent progenitors 3 (MPP3) while no other MPPs compartment was affected (**Figure 1B**). MPP3 are a myeloid-biased subset that constitutes a reservoir for granulocyte/macrophage progenitors ^23,24^ and are the only MPP subset to co-express CX3CR1 and CSF1R (**Figure 1C**). Consistent with a role for CUREA in early lineage commitment, *Csf1r* mRNA was significantly reduced in both MPP3 and in Lin^-^SCA^+^KIT^+^ (LK) cells in CUREA^-/-^ compared to WT (**Figure 1 D-E**). 5’RACE from sorted LK cells was consistent with initiation from the cluster of start sites (TSS) in the major myeloid promoter regardless of genotype (**Figure 1F).** There was no depletion of downstream progeny, granulocyte and macrophage progenitors (GMP), common myeloid progenitors (CMP), megakaryocyte and erythroid progenitors (MEP), megakaryocyte progenitors (MKP) (**Supp-Figure 2A-B**), or erythroid progenitors (**Supp-Figure 2C**) in CUREA^-/-^ mice. The changes in *Csf1r* mRNA were not reflected in CSF1R protein. As reported previously ^4^, the FRed reporter is first detected in GMP, but neither FRed nor CD115 was significantly altered in CUREA^-/-^ mice (**Supp-Figure 2D-F**).

**Figure 1.**
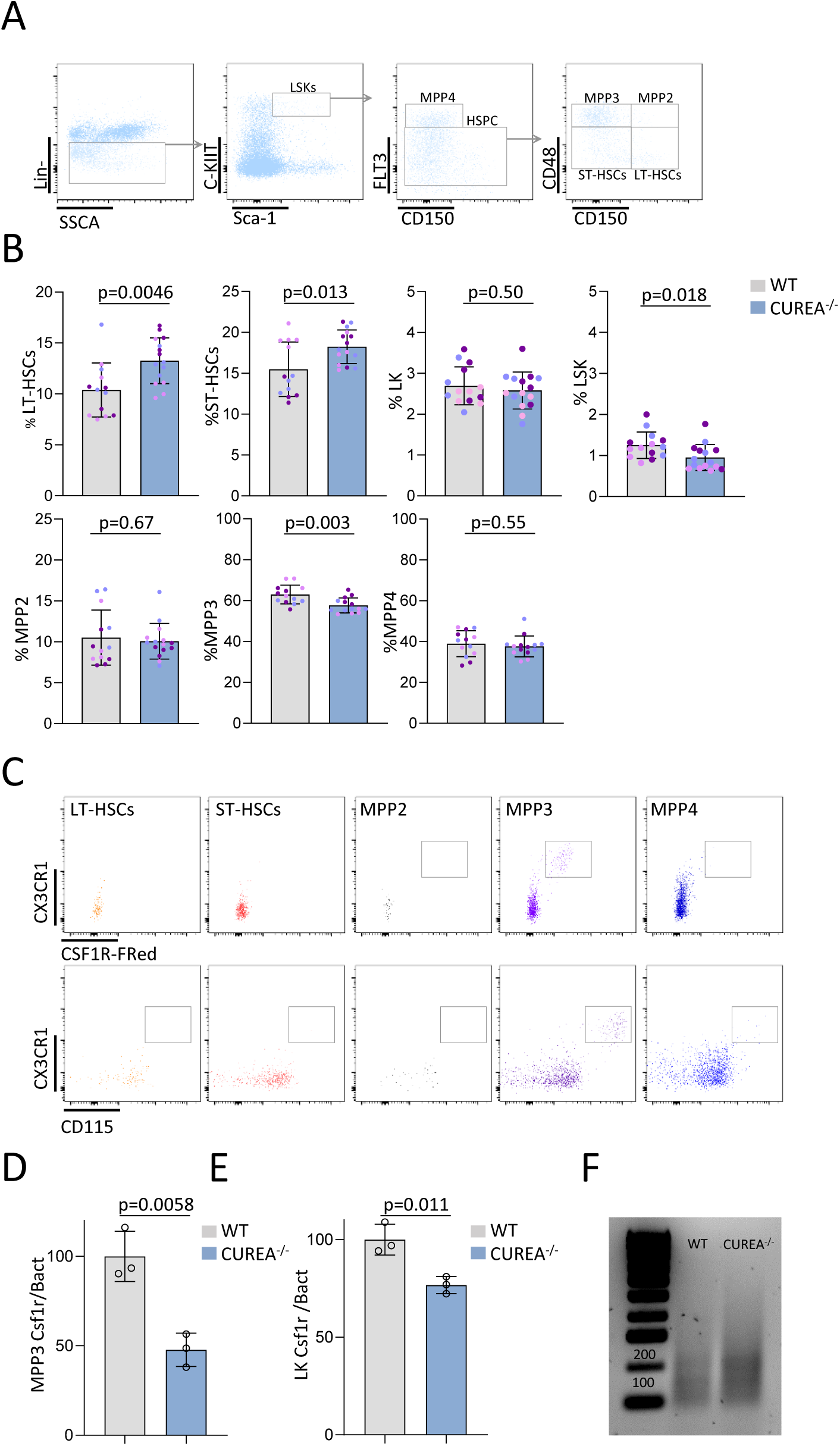
Hematopoietic differentiation in CUREA^-/-^ bone marrow. (A) Gating strategy for LKs, LSKs, MPPs, LT-HSCs, and ST-HSCs. (B) Proportions of HSPCs and myeloid progenitors in WT and CUREA^-/-^. Each dot represents a mouse, each color an experiment (pool of three experiments with at least four mice per group). (C) CX3CR1 and CSF1R expressions in HSPCs in CX3CR1-EGFP, CSF1R FRed mice. (D) Relative *Csf1r* mRNA expression in MMP3 and (E) LK of WT and CUREA^-/-^ mice. (F) *Csf1r* promoter capture (5’RACE) in LKs in WT and CUREA^-/-^ mice. Data show means and standard deviation. *p* values were determined by unpaired two-tailed t-test.

### CUREA is not required for myelomonocytic cell differentiation

Automated hematology analysis indicated no major effect on WBC composition in the blood, bone marrow and spleen of CUREA^-/-^ mice (**Figure 2A)**. Flow cytometry revealed a modest decrease in CD11b^+^, Ly6G^-ve^ cells in the blood and bone marrow but no change in the representation of classical (Ly6C^high^) monocytes and non-classical (Ly6C^low^) monocytes in any of the three organs investigated **(Figure 2B-C, Supp-Figure 3A-B)**. The FRed reporter provides a sensitive marker for monocytes that may be less sensitive to down-regulation by circulating CSF1 than CD115 ^25^. FRed was uniformly expressed in Ly6C^high^ monocytes. In the Ly6C^low^ population, MFI was around 4-5 fold higher but with a distribution tail suggesting a transition (**Figure 2D**). By contrast, CD115 was detected at similar levels in both populations. Classical monocytes can arise from monocyte-dendritic-cell progenitors (MDP-Mo) or granulocyte-monocyte derived progenitors (GMP-Mo) ^26,27^ and can be distinguished using CD319 and CD177 ^28^. GMP-Mo representation was marginally decreased in the blood of CUREA^-/-^ mice (**Figure 2B-C)**. *Csf1r* mRNA was reduced in the BM Ly6C^high^ monocyte and total splenocytes **(Supp-Figure 3C-D)** but neither the FRed reporter nor CD115 was altered in monocytes from the blood, BM or spleen (**Figure 2D Supp-Figure 3E-F).** To identify transcription start sites (TSS) in spleen, we performed 5’RACE from total splenocyte mRNA. The pattern was identical in both genotypes consistent with initiation in the major myeloid TSS (**Supp-Figure 3F)**.

**Figure 2.**
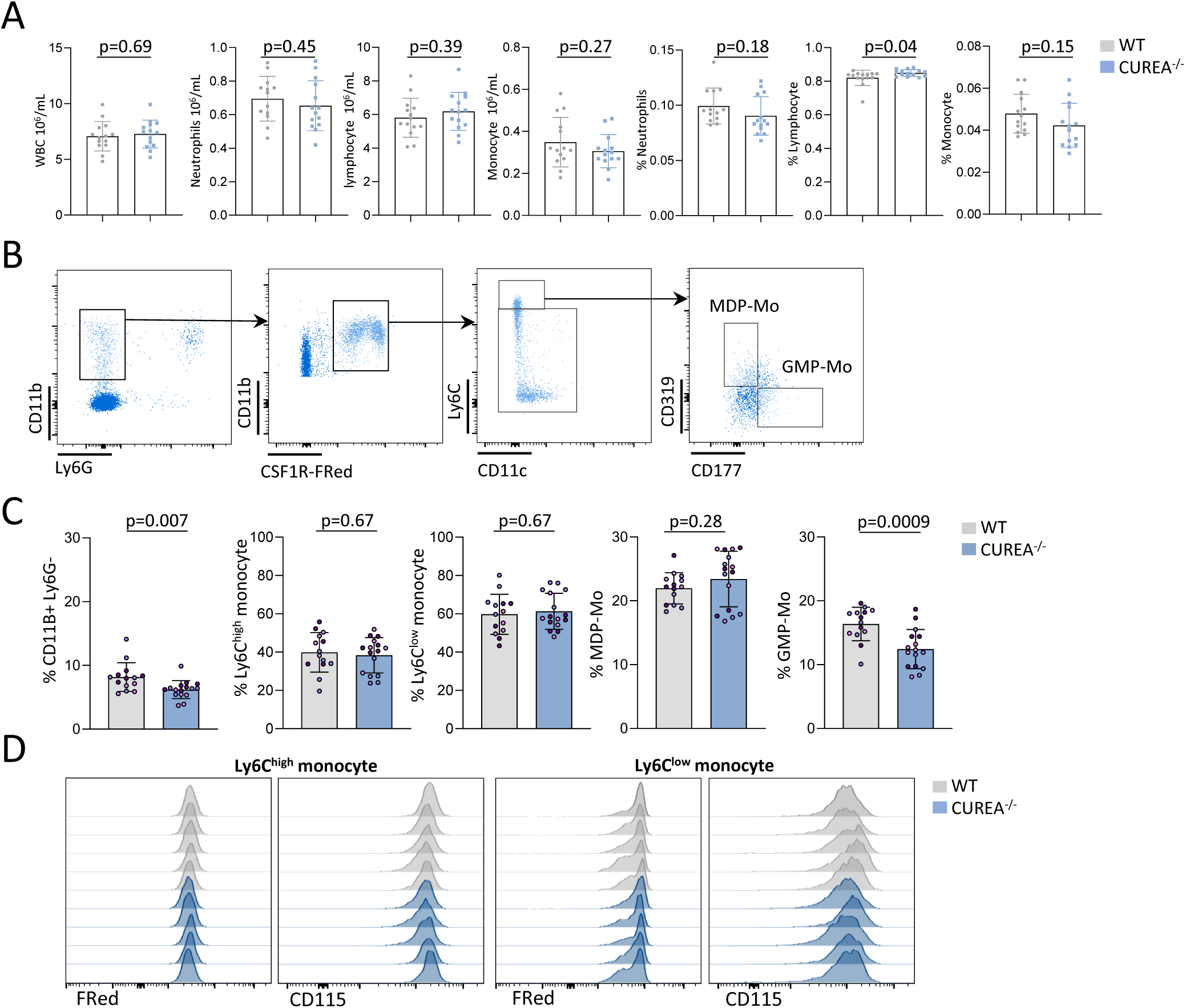
CUREA is not required for myelomonocytic differentiation. (A) Concentration and proportion of WBC, neutrophils, lymphocytes and monocytes within blood cells determined using a haematology analyser. (B) Representative gating strategy for blood monocytes and monocyte progenitor (GMP-MO, MDP-MO) populations. (C) Proportion of blood CD11b^+^, Ly6G^-ve^ (within single cells), Ly6C^high^, Ly6C^low^ monocytes (within CD11b^+^, Ly6G^-ve^, FRed^+^ cells), GMP-Mo, MDP-Mo (within Ly6C^high^ CD11c^-ve^ monocytes) in WT and CUREA^-/-^ mice. Each dot represents a mouse, each color an experiment (pool of three experiments with at least four mice per genotype). (D) FRed and CD115 MFI within Ly6C^high^ and Ly6C^low^ monocytes in WT and CUREA^-/-^ mice. Data show means and standard deviation. *p* values were determined by unpaired two-tailed t-test.

### CUREA does not impact stress hematopoiesis

The MacBlue transgenic mice were previously used to visualise monocyte reconstitution kinetics following cyclophosphamide administration in mice ^29^. Early studies on the mouse *Csf1r* promoter indicated that CUREA contains growth factor responsive elements ^30,31^ which might contribute to increased monocyte production in response to demand. Mice were injected i.p. with 200mg/kg of cyclophosphamide and monocyte mobilization assessed 5 days post treatment as described ^29^. Monocyte mobilisation was indistinguishable between WT and CUREA^-/-^ mice **(Supp-Figure3 G-I)**. We next investigated whether CUREA deletion could alter the monocytosis induced by CSF1 *in vivo.* Mice were injected with CSF1-Fc (1mg/kg) subcutaneously and BM, blood and spleen were collected at 3 and 5 days post treatment. As described previously ^32^ we observed thrombocytopenia, increased spleen weight (**Supp-Figure 4A-B)** and increased Ly6C^high^ monocytes in blood and spleen but there was no difference between WT and CUREA^-/-^ mice (**Supp-Figure 4C-E).**

### Resident tissue macrophages are unaffected in CUREA^-/-^ mice

The *Csf1r-*FRed reporter provides a unique marker to visualise resident tissue macrophages ^4^ but was not previously analysed in detail by Flow Cytometry. Based upon analysis of MacBlue mice^19^, the CUREA mutation was predicted to impact CSF1R expression in tissue macrophages. **Figures 3A-F** show combined detection of FRed, CD45, CD11b and F4/80 in cell populations obtained by disaggregation of liver, peritoneum, lung, heart, white adipose tissue (WAT) and kidney. In the peritoneal cavity the expression of FRed distinguished F4/80^high^ large peritoneal macrophages from F4/80^low^ small peritoneal macrophages (**Figures 3B**). In each of the organs analysed the FRed marker defined apparent subpopulations of macrophages. FRed^high^ populations are dominant in peritoneum, kidney, heart and adipose that are selectively depleted of resident macrophages in *Csf1r*^ΔFIRE/ΔFIRE^ mice ^10^. We found no significant difference in distribution of markers or level of FRed expression in subsets between WT and CUREA^-/-^ mice. The macrophages of the liver, spleen and intestine are not affected in *Csf1r*^ΔFIRE/ΔFIRE^ mice but the latter are replaced continuously by monocytes and depleted by anti-CSF1R treatment ^33^. F4/80 immunolocalisation confirmed that resident macrophages in these organs as well as the pancreas were not affected in CUREA^-/-^ mice (**Figure 3G, I)**.

**Figure 3.**
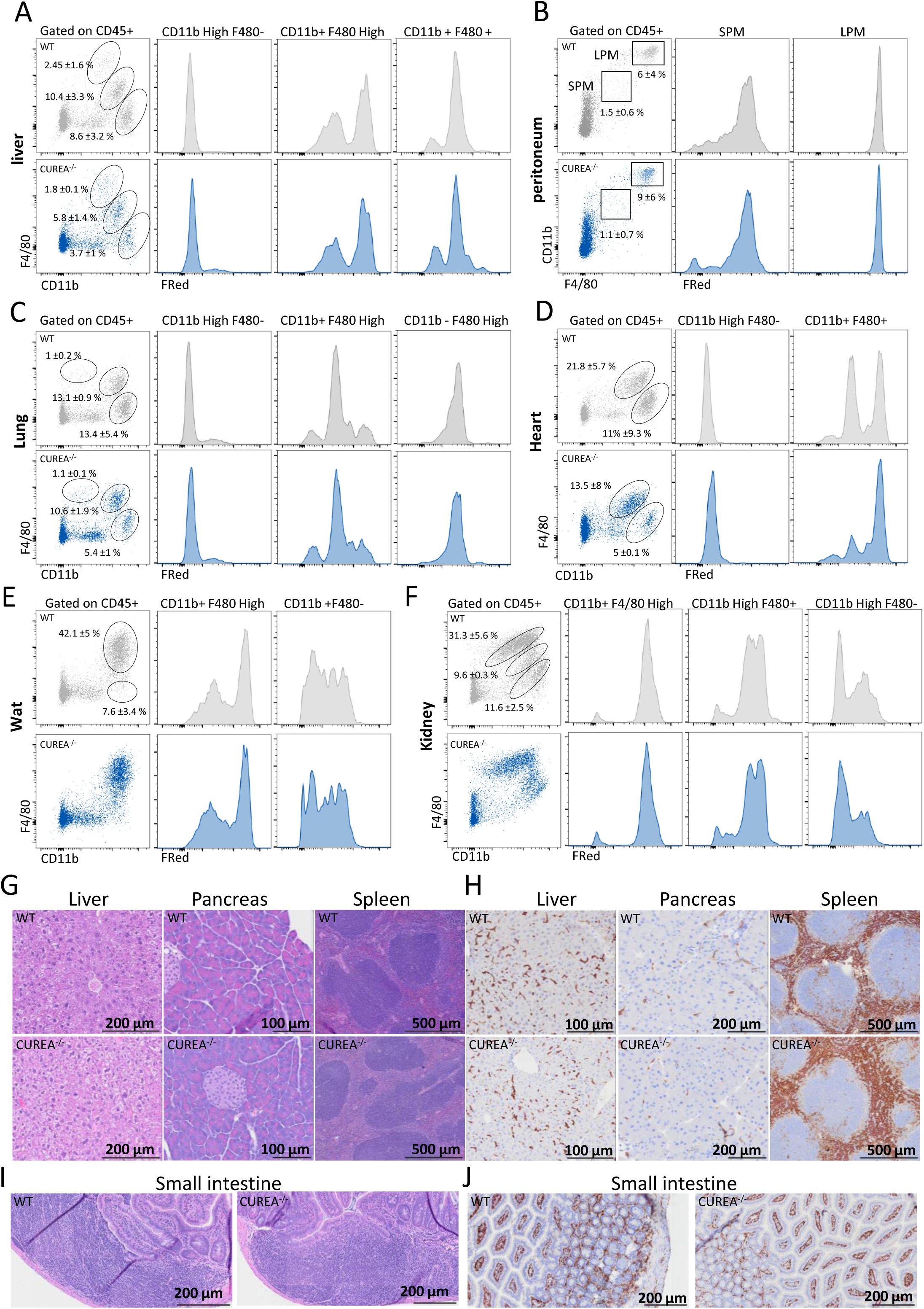
CUREA is not required for tissue resident macrophage differentiation. Representative gating strategy and proportion (within CD45+) of cell stained for CD45, CD11b, F4/80 and expressing Csf1r-Fred (shown with histogram) in (A) liver, (B) peritoneum showing small peritoneal macrophage (SPM) and large peritoneal macrophage (LPM), (C) lung, (D) heart, (E) white adipose tissue (WAT) and (F) kidney from adult WT and CUREA^-/-^ mice. Data show means and standard deviation. (G) Representative images of H&E and (H) F4/80 staining on liver, pancreas, spleen. (I) H&E and (H) F4/80 staining on small intestine from WT and CUREA^-/-^ mice.

### CUREA regulates the rate of *Csf1r* mRNA expression

We next investigated whether CUREA deletion alters the CSF1-dependent proliferation and differentiation of bone marrow-derived macrophages (BMDM). No change was observed in BMDM morphology and proliferation, CSF1R and FRed expression or *Csf1r* transcript level *in vitro* between WT and CUREA^-/-^ mice (**Figure 4A-C)**. CSF1R is continuously internalised and degraded in the presence of CSF1 ^34^. To monitor the contribution of CUREA to CSF1R turnover, we analysed a time course of CD115, FRed and *Csf1r* expression following CSF1 removal from CSF1-replete BMDM **(Figure 4E-F)**. Surface CSF1R (CD115) increased following removal of CSF1 and declined following CSF1 re-addition regardless of genotype. FRed also increased in response to CSF1 depletion, but was not acutely sensitive to CSF1 re-addition. However, there a significant delay in *Csf1r* transcript synthesis upon CSF1 starvation in CUREA^-/-^ BMDM compared to WT **(Figure 4G)**.

**Figure 4.**
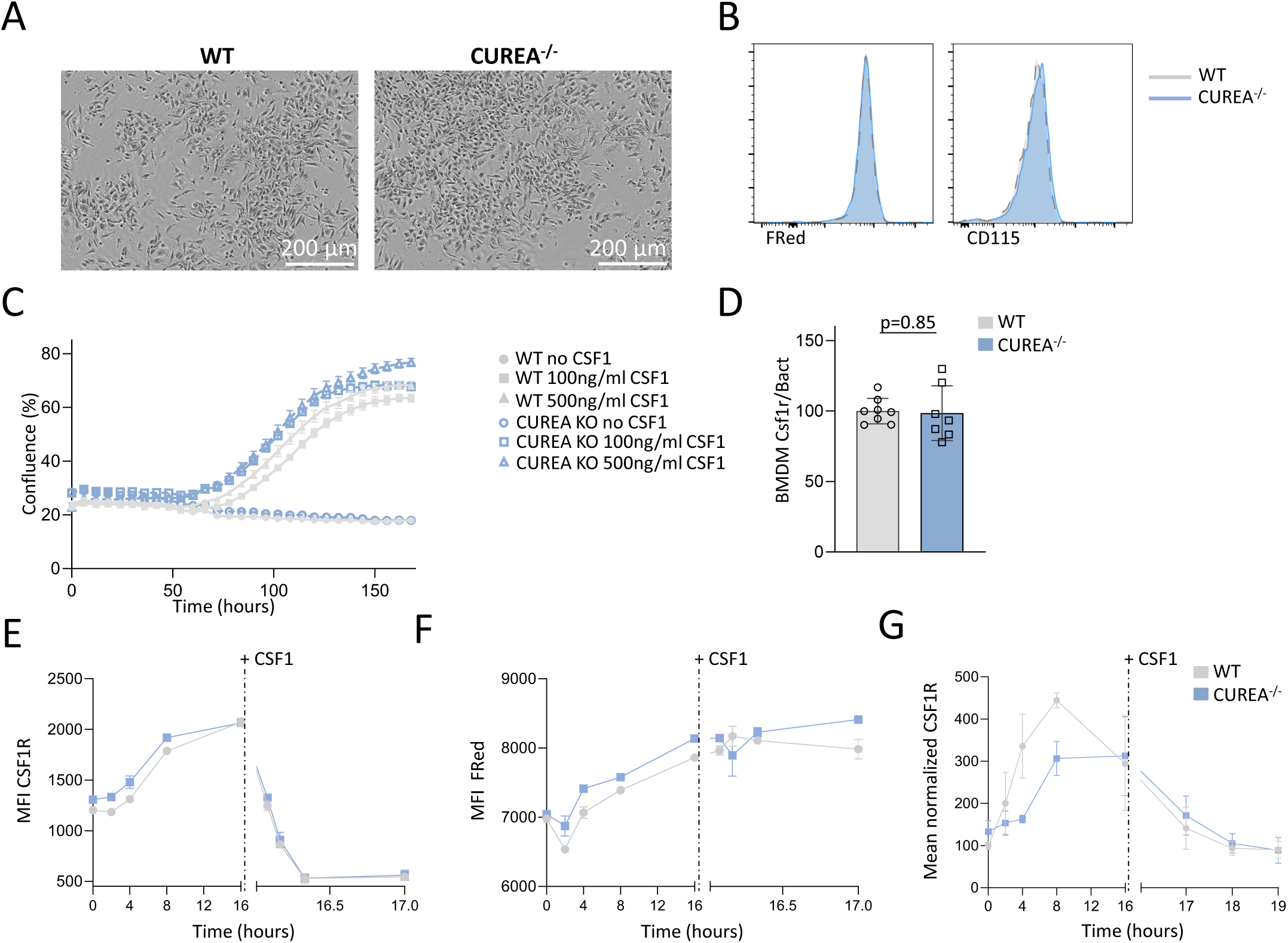
The effect of CUREA deletion on *Csf1r* transcription. (A) Representative image of WT and CUREA^-/-^ bone marrow derived macrophage (BMDM) cultured for 5 days in the presence of CSF1 (100ng/ml) (B) Representative flow cytometry detection of FRed and staining of CD115 in BMDM cultured for 7 days. (C) Bone marrow cells from WT or CUREA^-/-^ mice was pooled and cultured in 100 ng/ml or 500 ng/ml CSF1 for 7 days in an Incucyte with confluence measurement every 6 hours. Data show mean confluences and standard deviation for 6 wells per genotype per time point. (D) Relative *Csf1r* mRNA expression in BMDM from WT and CUREA^-/-^. (E and F) CSF1R internalization assay on WT and CUREA^-/-^ BMDM. Cells were washed and incubated without CSF1 starved for 16 hours, then restimulated for the indicated times. (E) CD115 and (F) FRed expression were assessed by flow cytometry at the specified time points. (G) Relative expression of *Csf1r* transcript in WT and CUREA^-/-^ BMDMs upon CSF1 starvation, followed by CSF1 restimulation. Data show means and standard deviation. *p* values were determined by unpaired two-tailed t-test. Reduced expression of CD115 in CUREA^-/-^ is significant at 4 and 8hrs post CSF1 removal (p<0.001).

### CUREA contributes to microglial homeostasis

The MacBlue transgene is expressed by microglia suggesting that CUREA is not absolutely required for expression of *Csf1r.* In flow cytometry performed on digested brain tissue, high expression of FRed distinguished CD45^low^/CD11b^+^ microglia from a more heterogeneous CD45^high^/CD11b^+^ population. CD45 is highly-expressed by brain-associated macrophages (BAM) in WT animals but also characterises monocyte-derived microglia-like cells and damage-associated microglia ^35^. There was no difference in CSF1R-FRed expression between WT and CUREA^-/-^ in either brain myeloid population but a significant decrease in microglia and increase in BAM **(Figure 5A-B; Supp**-**Figure 6A**). Immunofluorescence staining of microglia in the brain cortex (IBA1^+^, CSF1R-FRED^+^) and perivascular cells in the meninges (IBA1^+^, CD169^+^) did not highlight any morphological or phenotypic changes in CUREA^-/-^ (**Figure 5C-D**). The absence of microglia in *Csf1r*^ΔFIRE/ΔFIRE^ mice is associated with extensive calcification of the thalamus ^36^ but no such pathology was detected in WT or CUREA^-/-^ mice at 37 weeks of age. Similarly, microglia are implicated in the regulation of perineuronal nets ^37^ but their detection using *Wisteria floribunda* agglutinin or localisation of parvalbumin interneurons staining was not affected (**Supp-Figure 6B-D).**

**Figure 5.**
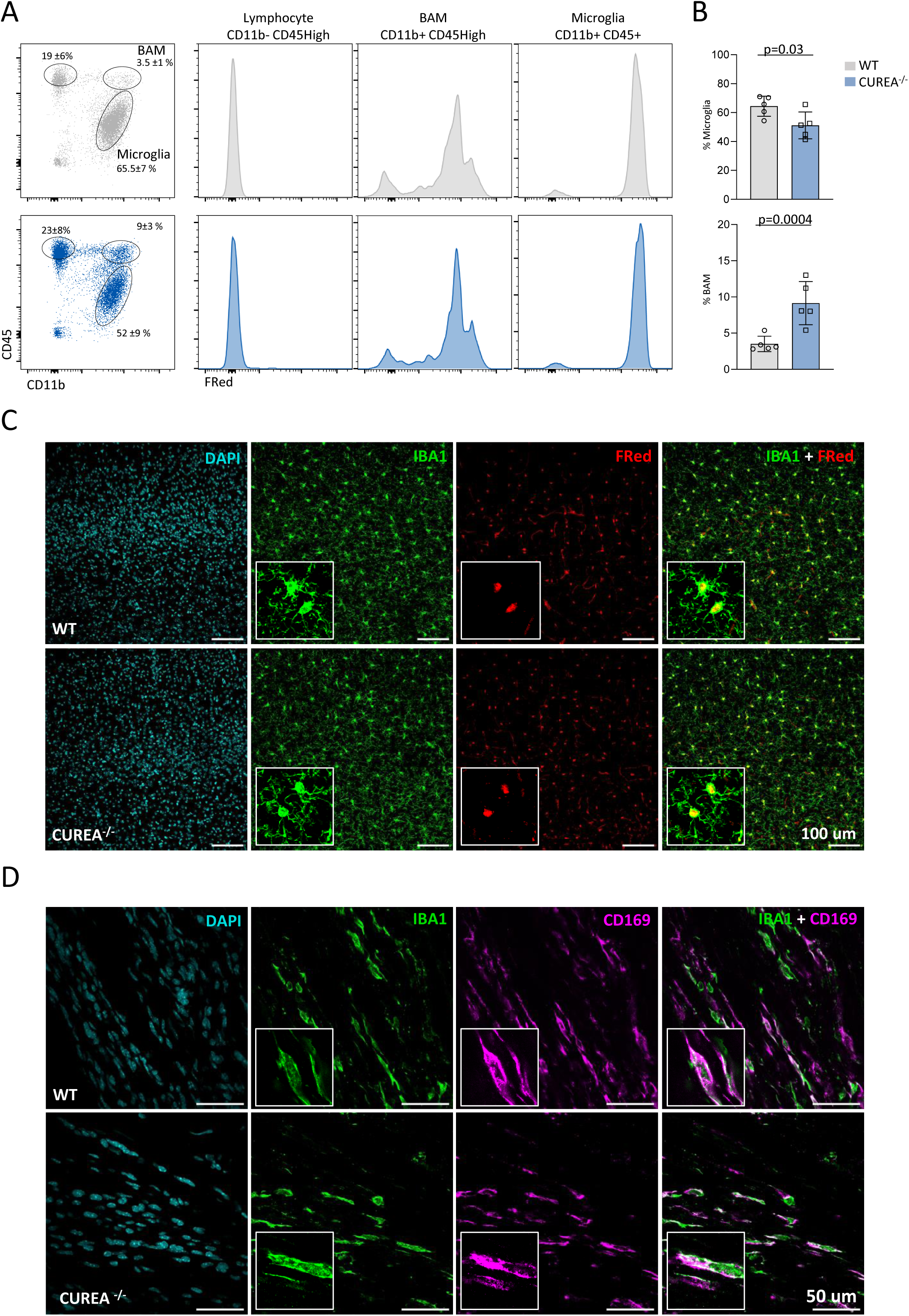
CUREA deletion impacts microglial phenotype. (A) Gating and FRed expression for microglia and brain-associated macrophages (BAM) from disaggregated brain. Cells were stained for CD45 and CD11b and analysed by flow cytometry. (B) Proportion of microglia and BAM in WT and CUREA^-/-^ brains. Each point is an individual animal. (C) Free floating sections of WT and CUREA^-/-^ cortex were stained to detect IBA1 with *Csf1r*-FRed expression. Inset panels highlight colocalization between IBA1 and FRed. (D) Isolated meningeal layers of WT and CUREA^-/-^ brain stained for IBA1 and CD169. Inset panels highlight colocalization of the two markers.

### CUREA regulates osteoclast differentiation and function

In addition to trophoblast promoter activity, CUREA contains a promoter utilized by osteoclasts (OCL) and deletion of the element in a multicopy *Csf1r*-EGFP transgene impacted OCL-specific reporter gene expression^17,18^. We first investigated whether *in vivo* osteoclast distribution was dependent upon CUREA. As reported previously^4^, *Csf1r-*FRed was highly-expressed by OCL on the bone surface, predominantly visible around the bone head, and colocalised with tartrate-resistant acid phosphatase (TRAP). At a qualitative level there was no detectable difference between WT and CUREA^-/-^ mice in terms of OCL location or abundance or FRed fluorescence intensity **(Figure 6A).** However, micro-CT scans on the femurs of 12-week-old male mice revealed a significant increase in trabecular bone complexity in CUREA^-/-^ mice **(Figure 6B, C; Supp-Video 1-2).** To determine the possible mechanism, we assessed osteoclast differentiation *in vitro* using CSF1 and RANKL as described ^17^ (**Figure 6D**). We observed delayed differentiation of mature TRAP^+^ osteoclasts **(Figure 6E)** and decreased *Csf1r* transcript levels in CUREA^-/-^ cultures on day 7 **(Figure 6F**). To confirm the OCL-specific TSS described previously we performed 5’RACE. We detected an OCL-specific *Csf1r* transcription start site (TSS) in WT that was greatly reduced in CUREA^-/-^ cultures **(Figure 6G)**. We the performed nested PCR using a primer specific for the osteoclast transcript (F1.2). The primer sequence is immediately 3’ of the CUREA deletion. The locations of all of the regulatory elements, primers, splice sites and CUREA deleted sequence are shown in **Figure S7**. We detected a PCR product in WT OCL, but not in spleen and LK bone marrow cells **(Figure 6H).** This band was not present in CUREA^-/-^ cultures.

**Figure 6.**
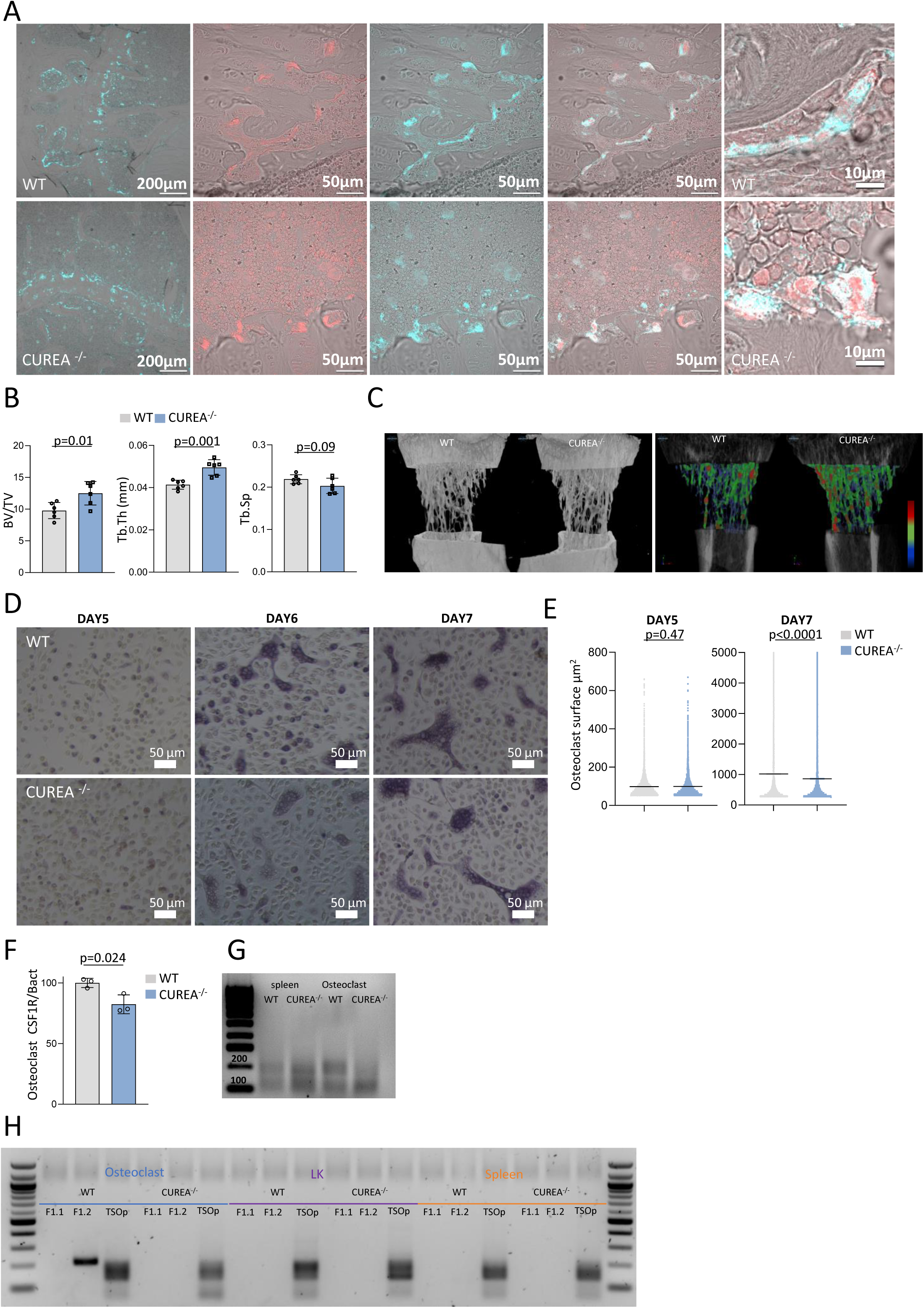
CUREA regulates *Csf1r* expression and osteoclast function. (A) Femoral sections from adult 22-week-old WT and CUREA^-/-^ mice stained for TRAP as described in Materials and Method. Representative images show the colocalisation of TRAP (cyan) and FRed at different magnifications (B) Quantification of bone volume/trabecular volume (BV/TV), trabecular thickness (Tb.Th), and trabecular space (Tb.Sp) from micro-CT imaging in adult male WT and CUREA^-/-^ mice. Data show means and standard deviation. *p* values were determined by unpaired two-tailed t-test. (C) Representative micro-CT 3D reconstruction of femur diaphysis and trabecular bone from WT and CUREA^-/-^ mice. Panel at right shows clusters of bone density (red high density, blue low density) within trabecular bone in the two genotypes. (D) Bone marrow cells from WT and CUREA^-/-^ mice were cultured in 100ng/ml CSF1 for 3 days, 40ng/ml RANKL was then added, the cells cultured until day 5, 6 or 7 and then stained for TRAP. Panels show representative images at each time point. (E) Surface area of individual osteoclasts in bone marrow cultures from WT and CUREA^-/-^ was analysed as described in Materials and methods. Each point represents a single TRAP^+^ OCL. Data show means with standard error of the mean (SEM). *p* values were determined by Man Whitney unpaired two-tailed t-test. (F) Relative *Csf1r* mRNA expression in WT and CUREA^-/-^ osteoclast cultures after 7 days (G) 5’RACE to detect *Csf1r* transcription starts in mRNA isolated from splenocytes and osteoclasts of WT and CUREA^-/-^ mice. (H) Nested PCR analysis of products of 5’RACE using the F1.2 primer to detect OCL-specific transcripts. Note the OCL-specific amplicon in WT bone marrow cultures that is absent in spleen and progenitor cells and in CUREA^-/-^ mice.

### CSF1R is expressed in trophoblast cells

*Csf1r* mRNA is highly expressed expressed in trophoblasts ^8^ but protein expression has not been previously confirmed or localized. *Csf1r*-FRed has also not previously been used to visualize macrophages in the embryo. We performed time matings and assessed CSF1R-FRed expression within the embryo at day 11.5 of development (E11.5), using tissue clearing. Diffuse FRed^+^ macrophage distribution was observed throughout the embryo with more obvious concentrations in the head and eyes as well as the liver of both WT and CUREA^-/-^ embryos (**Figure 7A)**. In the placenta, FRed detected by anti-RFP was highly expressed in TROMA1^+^ trophoblasts, notably trophoblast giant cells. Placentas from WT and CUREA^-/-^ embryos were indistinguishable in terms of FRed expression, overall morphology **(Figure 7B)** or the level of *Csf1r* mRNA **(Figure 7C).** To understand how *Csf1r* is expressed despite the deletion of the trophoblast-specific promoter we performed 5’RACE. In WT placenta we detected two bands corresponding to the major TSS identified previously and two alternate splice acceptors ^8^. In CUREA^-/-^ placenta, the two bands were still present but both displaced to somewhat higher size **(Figure 7D)**. Nested PCR using a primer (F1.1) based on the known TSS within CUREA confirmed that these two bands are indeed the known splice variants of the WT trophoblast transcript. As predicted, they were not detected in the mutant placenta. However, primer F1.2, immediately downstream of CUREA but 5’ of the predicted splice donor yielded two bands in both WT and CUREA^-/-^ placenta. We conclude that in the mutant transcripts are initiated from TSS upstream of CUREA but retain the same splicing pattern.

**Figure 7.**
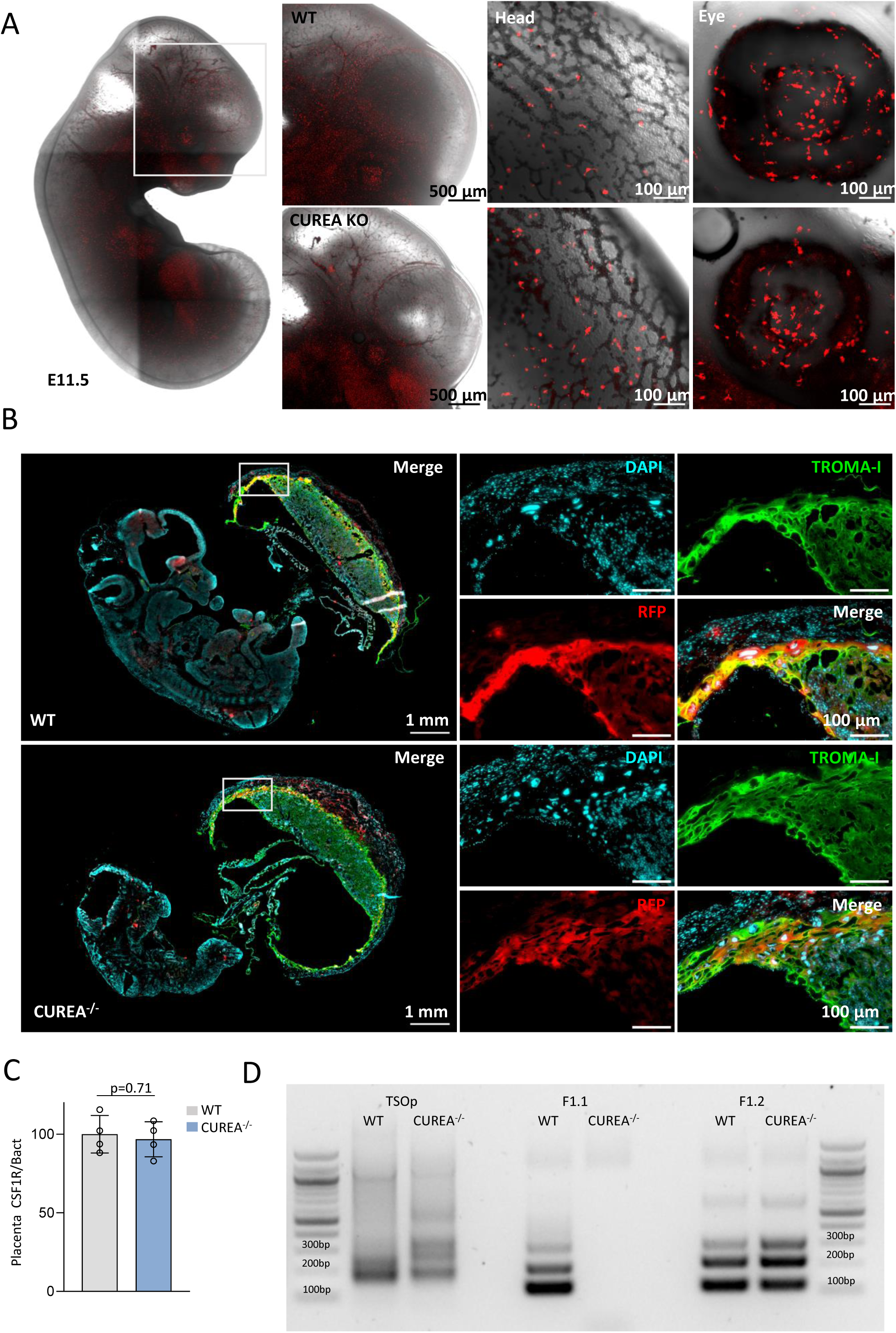
Expression of CSF1R in trophoblast cells. (A) Whole mount confocal imaging of tissue cleared embryo (E11.5) showing FRed expressing cells. Highlighted region shows macrophages within the head and eyes of WT and CUREA^-/-^ embryo (B) WT and CUREA^-/-^ embryos (E11.5) with placenta sectioned and stained for TROMA-I, RFP, and DAPI. (C) Relative *Csf1r* mRNA expression in placenta from WT and CUREA^-/-^ embryos Data show means and standard deviation. *p* value was determined by unpaired two-tailed t-test. (D) 5’RACE analysis of transcription initiation in placenta of WT and CUREA^-/-^ E11.5 embryos. Nested PCR was performed using *Csf1r-*specific reverse primer (R2) and forward primers TSO, F1.1, or F1.2 (see Figure S7) as indicated.

## Discussion

### Partial redundancy of *Csf1r* enhancer elements

In the study we took the novel approach of deleting a candidate enhancer element on the background of an integrated *Csf1r-*FusionRed reporter. We focused on the CUREA element because previous studies had shown that this element was required for expression of a multi-copy transgene in most resident tissue macrophages ^17,19^. However, as in our previous analysis of deletion of the intronic enhancer, FIRE^10^, we found that the effect of genomic deletion of the CUREA element on macrophage development did not correspond to the effect on transgene expression. In the specific case of liver Kupffer cells (KC), which do not express the MacBlue transgene ^19^ or multicopy *Csf1r-* EGFP transgenes lacking the FIRE sequence ^17^, chromatin analysis of mouse KC ^13^ suggests the explanation. The ATAC-seq data in this study indicates that both FIRE and CUREA are in open chromatin in KC but at least 4 additional candidate enhancers lie outside of the 7.2kb mouse *Csf1r* genomic sequence (3.5kb promoter plus first intron) that is sufficient to produce robust lineage-specific reporter gene expression in KC in multiple species ^2,38–40^. Datasets from the Glass lab^13,15,16^ and others (reviewed in ^9^} confirm the activity of multiple enhancers in other tissue macrophage populations including microglia and their occupation by lineage specific transcription factors, notably PU.1.

Our use of the *Csf1r-*FusionRed reporter provided no evidence of a significant effect of CUREA deletion. However, it did extend the analysis of the utility of this tool. We confirmed for the first time that trophoblast expression of *Csf1r* mRNA^8^ is translated to protein. The role of CSF1R in placenta remains unclear since placental and embryo development appears unaffected in *Csf1r*^-/-^ rats and mice ^1^. As shown in Figures 2, 3 and 4, *Csf1r-*FRed expression distinguishes Ly6C^high^ and Ly6C^low^ monocytes and heterogeneity amongst F4/80^high^ tissue-resident macrophages. The highest levels of FRed were detected in peritoneal macrophages, microglia and kidney and heart F4/80^high^ macrophages, each selectively depleted in *Csf1r*^ΔFIRE/ΔFIRE^ mice ^10^.

### CUREA is not entirely redundant

Since the MacBlue transgene is highly-expressed by monocytes ^19^ and CSF1R is not absolutely required for monocytopoiesis^33^ we did not anticipate a major effect of CUREA deletion on monocytic differentiation. However, the promoter and CUREA are in open chromatin in myeloid progenitors prior to chromatin remodelling of FIRE^11^. Consistent with a functional role of CUREA in early lineage commitment we found a small but reproducible accumulation of LT-HSCs and ST-HSCs associated with decreased MPP3 representation within CUREA^-/-^ bone marrow and significant reduction in *Csf1r* mRNA in both populations. The CUREA deletion also reduced the rate of re-expression of *Csf1r* following removal of CSF1 from BMDM. However, these differences were not sufficient to alter myeloid cell mobilization in response to cyclophosphamide or CSF1-Fc.

The only tissue-resident macrophage population that was impacted measurably by CUREA deletion was the microglial population of the brain. We observed a polarization of the resident population towards the phenotype (CD45^high^/CD11b^low^) associated with so-called brain associated macrophages and monocyte-derived microglial replacement ^41^. In WT mice there is a significant population of microglia that lacks expression of the MacBlue transgene^19,21^. We suggest that the CUREA mutation interferes with CSF1R-dependent microglial homeostasis leading to an acceleration of replacement by monocyte-derived cells with age^41^. Interestingly, ChIP-seq analysis of mouse microglia ^15^ indicates that key transcription factors SALL1 and SMAD4 that control homeostatic microglial identity are both bound to CUREA.

CUREA contains a promoter utilized specifically by bone-resorbing OCL^17^. We confirmed the existence of OCL-specific TSS in WT OCL cultures and their absence in CUREA^-/-^ OCL (**Figure 6H, I**). CUREA deletion led to a small decline in *Csf1r* mRNA in culture, delayed OCL differentiation and a significantly increased trabecular complexity *in vivo*. However, the effect is not comparable to the complete OCL deficiency and severe osteopetrosis seen in *Csf1r-/-* mice ^22^. It appears that expression from the downstream major myeloid TSS is sufficient to maintain *Csf1r* expression in OCL.

Finally, we considered expression of *Csf1r* in trophoblasts. CUREA contains TSS expressed only in placenta and transcripts initiating from the major myeloid promoter were almost undetectable in placenta ^42^. The predominant trophoblast TSS is associated with a TATA box (**Figure S7**). Surprisingly, CUREA deletion had no effect on trophoblast expression of *Csf1r* mRNA or the FRed reporter. In this case, the TSS apparently moved upstream to a cryptic promoter that was sufficiently active to maintain expression at the same level. Although the CUREA sequence is conserved between mouse and human, the TATA box and transcription initiation at this location in trophoblasts is not. In humans and other large animals, *CSF1R* transcription initiates 20kb upstream in the 3’ end of the *PDGFRB* gene^7^. The CUREA element is approximately the same size as a nucleosome. It may be that the position of the TSS is determined by nucleosome phasing ^43^ rather than the precise sequence and is simply displaced upstream by the removal of 176bp.

## Conclusion

Enhancer redundancy is a common feature of regulated gene expression in differentiation and development^44,45^. Some elements appear more essential than others. Germ-line deletion of FIRE has a profound effect on *Csf1r* transcription and differentiation of specific macrophage populations but is not absolutely required for *Csf1r* expression^10^. Similar penetrant impacts on lineage-specific expression have been observed in mice with deletions of myeloid-specific enhancers in other loci ^46,47^. By contrast, the effects of CUREA deletion are subtle; more typical of the partial redundancy of so-called shadow enhancers^44,45^. It is possible that CUREA interacts in *cis* with other elements including FIRE, as seen with enhancers in the *Irf8* locus^48^. Nevertheless, the previous studies with the transgenic reporters and ChIP-seq analysis of binding of multiple transcription factors indicate that CUREA function distinguishes monocytes from tissue resident macrophages. Our analysis has identified non-redundant functions of CUREA in *Csf1r* transcription. Combinatorial deletion with other conserved elements will be required to establish the interaction with other candidate enhancers, including FIRE, that function in tissue-specific *Csf1r* expression.

## Acknowledgements

The authors are grateful for the discussions and helpful advice from Anthony Adamson. The generation of the CUREA mice was funded originally by a grant from the Medical Research Council (MRC) UK grant MR/M019969/1 to DAH. This work was supported by core support and direct funding from The Mater Foundation and NHMRC Investigator Grant #2007850 to DAH. We acknowledge input and expertise from the Biological Resources Facility and the Preclinical Imaging, Microscopy, Histology and Flow Cytometry facilities of the Translational Research Institute (TRI). TRI is supported by the Australian Government.

## Competing Interests

The authors declare no competing interests

---

**Supplementary Figure 1. Generation of *Csf1r T2A-FRed* ^ΔCUREA/ΔCUREA^ (CUREA^-/-^) mutant mice**

(A) Schema of the Csf1r chromatin landscape, CRISPR deletion strategy for CUREA deletion. (B) CUREA^-/-^ deletion following CRISPR cut and (C) genotyping from deleted region flanks. (D) Sequences from wild type (WT) and selected edited patterns from CUREA^-/-^

**Supplementary Figure 2. CUREA regulation of myeloid differentiation**

(A) Proportion of GMP, CMP, MEP within LKs. (B) Proportion of MKPs within LKs. Each point represents a mouse, each color an experiment (pool of two experiments with at least four mice per genotype). (C) Representative flow cytometry gating strategy for the erythroid progenitor cell population and percentage of erythroid progenitor populations in total bone marrow cells. (D) CD115 mean fluorescent intensity (MFI) within LKs, LSKs, MPPs, LT-HSCs, and ST-HSCs. (E) CD115 MFI of GMP, CMP, MEP in WT and CUREA^-/-^.Each dot represents a mouse, each color an experiment (pool of two experiments with at least three mice per group). Data show means and standard deviation. *p* values were determined by unpaired two-tailed t-test. (F) FRed MFI within LT-HSCs, ST-HSCs, MPPs, LSKs, LKs, GMP, CMP, MEP in WT and CUREA^-/-^ mice. Each histogram represents a mouse. Data show means and standard deviation. *p* values were determined by unpaired two-tailed t-test.

**Supplementary Figure 3. Analysis of committed progenitors and mature myeloid cells**

(A) Proportion of bone marrow and (B) spleen CD11b^+^, Ly6G^-ve^ (within single cells), Ly6C^high^, Ly6C^low^ monocytes (within CD11b^+^, Ly6G^-ve^, FRed+ cells), GMP-Mo, MDP-Mo (within Ly6C^high^ CD11c^-ve^ monocytes) in WT and CUREA^-/-^ mice. Each point represents a mouse, each color an experiment (pool of two or three experiments with at least 3 mice per group). (C) *Csf1r* mRNA expression in Ly6C^high^ monocytes and (D) splenocytes. (E) FRed and CD115 MFI in bone marrow and spleen Ly6C^high^ and Ly6C^low^ monocytes from WT and CUREA^-/-^ mice. Each histogram represents a mouse. (F) Identification of *Csf1r* TSS by 5’RACE on splenocyte mRNA from WT and CUREA^-/-^. (G) Proportion of BM (H), blood and (I) spleen CD11b^+^, Ly6G^-ve^ cells, neutrophils (within single cell), Ly6C^high^ and Ly6C^low^ monocytes (within CD11b^+^, Ly6G^-ve^, FRed+ cells) in non-treated (NT) or cyclophosphamide-treated WT and CUREA^-/-^ mice harvested at day 5. Data show means and standard deviation. *p* values were determined by unpaired two-tailed t-test.

**Supplementary Figure 4. Analysis of response to CSF1-Fc in WT and CUREA^-/-^ mice.**

Mice were treated with a single injection of 5mg/kg CSF1-Fc subcutaneously and analysed 3 and 5 days later. (A) Platelet numbers determined using haematology analyser. (B) Spleen weight following CSF1 administration. (C-E) Proportions of neutrophils (Ly6G^+^), monocytes (CD11b^+^Ly6G^-ve^FRed^+^ and Ly6C^high^ and Ly6C^low^ monocytes (within monocyte population) in marrow (C), blood (D) and spleen (E). Data show means and standard deviation. *p* values were determined by unpaired two-tailed t-test.

**Supplementary Figure 5. Localization of CSF1R-dependent tissue-resident macrophages**

Sections of (A) liver, (B) pancreas, (C) spleen of adult WT and CUREA^-/-^ mice were stained for DAPI, IBA1 and CD169 as indicated.

**Supplementary Figure 6. Absence of brain pathology in CUREA^-/-^ mice.**

(A) Gating strategy for microglia and BAM. (B) Images show the thalamus from 30 week old WT and CUREA^-/-^ mice stained with risedronate (RIS; purple), and DAPI (blue). For comparison, image at right shows extensive calcification detected in the thalamus of microglia-deficient mice at similar age ^49^ (C) IF images of cortex from WT and CUREA^-/-^ mice showing perineuronal nets (WFA), parvalbumin interneurons (PV), FRed expression, and DAPI. (D) Percentage area of WFA and PV staining in somatosensory cortex quantified using ImageJ Fiji. Graphs show mean ± SEM from three mice per genotype, p values were determined using an unpaired two-tailed t-test with Welch’s correction.

**Supplementary Figure 7. Regulatory elements in the mouse *Csf1r* promoter**

## Notes

### Competing Interest Statement

The authors have declared no competing interest.

