## Supplementary material and figures for "A conserved upstream element in the mouse *Csf1r* locus contributes to transcription in hematopoietic and trophoblast cells"

Maxwell et *al*

#### Supplementary method

##### qPCR

RNA was extracted from cells using the PicoPure RNA Isolation Kit (Thermo Fisher Scientific) and cDNA synthesis with the SensiFAST cDNA Synthesis Kit (Meridian Bioscience), both according to the manufacturer's instructions. qPCR was performed using SYBR Green PCR Mater Mix (Thermo Fisher Scientific) on an Applied Biosystems QuantStudio 7 Flex system. Cycling conditions: 50°C 2 min, 95°C 10 min, followed by 40 cycles at 95°C 15 sec, 60°C 1 min, followed by 95°C for 15 sec, 60°C for 1 min and 95°C for 15 sec. Primers: *Csf1r* 5' CTCTTCACTCCGGTGGTGGT 3', 5' ACCTGGTACTTCGGCTTCTGC 3', *Actb* 5' CTCTGGCTCCTAGCACCATGAAGA 3', 5' GTAAAACGCAGCTCAGTAACAGTCCG 3'.

##### Blood analysis

Peripheral blood was collected into EDTA-coated tubes (Greiner K3, Interpath, Australia) and analysed using a Mindray (BC-5000) haematological analyser. Peripheral blood smears were stained with Wright-Giemsa (BIO Scientific, Australia).

##### Cell isolation and culture

Bone marrow was flushed from long bones by syringe with 2% fetal bovine serum (FBS) in phosphate buffered saline (PBS). Bone marrow cytospins were generated from $5x10^4$ cells, stained with Wright-Giemsa and imaged using a Nikon 50i microscope. To generate bone marrow-derived macrophages (BMM), ca. $10^7$ cells were seeded on 100mm square bacteriological plastic (Sterilin,

Thermo-Fisher, Australia) for bulk culture in 25 ml of complete medium (RPMI + 10% FBS, 25 U/mL penicillin, and 25μg/mL streptomycin (Gibco, Thermo-Fisher, Australia) and differentiated for 7 days with the addition of recombinant CSF1-Fc [1] (100ng/ml). For monitored differentiation assays $5x10^4$ bone marrow cells were seeded in 96 well coated plates in complete medium supplemented with 100 ng/ml CSF1-Fc. Cell confluence was monitored every 6 h for 7 days using the Incucyte (Sartorius, Australia) cell-by-cell analysis system.

For osteoclast culture bone marrow cells were harvested as described above and seeded at $5 \times 10^4$ cells/ml in 12 well tissue culture plates in complete medium with CSF1 (100 ng/ml) for 3 days. From day 3 the medium was replaced daily with medium containing CSF1-Fc (100 ng/ml) and RANKL (R&D Systems, 40 ng/ml) until day 7. Tartrate-resistant acid phosphatase (TRAP) staining was performed in situ using a kit (Cat #387A, Sigma, Australia) according to the manufacturer's instructions.

Peritoneal cells were obtained by flushing the peritoneal cavity with 10 ml PBS and prepared for flow cytometry staining by centrifugation at 400×g for 5 min at 4 °C prior to resuspension in FACS Buffer (FB, PBS containing 2% endotoxin-free FBS). Splenocytes were isolated by gently dissociating tissue with a 5 ml syringe in PBS, passed through a 40 μm filter, pelleted, subjected to red cell lysis (PharmLyse™, BD Biosciences, Australia), washed and resuspended in FB.

**Tissue disaggregation and flow cytometry.**

For cell isolation, tissues were dissected, minced with scissors in 10 ml collagenase solution (Hank's balanced salt solution containing 20 μl 10 mg/ml DNase1 (Roche), 10 mg collagenase IV and 1 mg dispase (Life Technologies)) and incubated in a shaking incubator at 37 ºC for 45 min. After incubation, tissues were mashed through a 70μm cell strainer, suspended in FB and centrifuged at 400 g for 5 min at 4 ºC. For lung, heart and kidney cells the supernatant was removed, the cell pellet was resuspended in 2 ml PharmaLyse and incubated at 4ºC for 10 min. The lysis reaction was then quenched with 10 ml FB, the cells centrifuged at 400 g for 5 min at 4ºC then resuspended in 5 ml FB. For brain and liver cells the pellet was resuspended in 12.5 ml isotonic Percoll (4.22 ml Percoll (GE Healthcare, Australia), 0.47 ml 10x PBS, 7.82 ml 1x PBS). Cells were centrifuged at 800 g 4ºC for 30 min with low acceleration and no brake. The myelin layer and supernatant was removed the cell pellet resuspended in 2 ml PharmaLyse and incubated on ice for 10 min. For adipose, cells were centrifuged at 500 g for 10 min with low acceleration and no break beginning at room temperature and cooling to 4 ºC. The adipose layer and supernatant was removed, the cell pellet resuspended in 2 ml PharmaLyse and incubated on ice for 10 min. Cells were diluted with FB, centrifuged and resuspended in FB for flow cytometry. For cell cycle analysis, sales were incubated in Hoechst 33342 (BD, 5mg/ml in PBS) for 5-10 mins then washed 3x in PBS.

Single-cell suspensions from blood, bone marrow and splenocytes were stained with fluorophore-conjugated antibodies on ice and analysed on an LSR Fortessa, (BD, Australia). Cell-surface markers used to define hematopoietic cell types and antibodies are provided in Supplemental Table 1. Data were analysed using FlowJo (BD).

**Immunofluorescence staining**

Brains were dissected from the skull and immersion fixed in 4% buffered formaldehyde for 5 h at room temperature (RT), washed once in PBS, and stored in PBS with 0.1% sodium azide. Brains were serially sectioned at 30 μm in the sagittal plane using a Leica VT1200S vibratome. Sections were permeabilised (1% Triton-X100, 0.1% Tween-20 in PBS) for 1 h at RT, blocked (4% serum, 0.3% Triton-X100, 0.05% Tween-20 in PBS) for 1 h at RT, and then stained with primary antibodies rabbit anti-IBA1 (1:500, Novachem, 019-19741) and guinea pig anti-parvalbumin (1:500, Synaptic Systems, 195 004) diluted in blocking buffer overnight at 4◦C. To label perineuronal nets, sections of somatosensory cortex were co-stained with *Wisteria floribunda* agglutinin (WFA; Merck, L1516) diluted 1:500 in blocking buffer. To label calcification, sections of thalamus were co-stained with AF647-RIS Imaging Reagent (BioVinc LLC, BV500101) diluted 1:500 in blocking buffer. Sections were then washed 3 x 5 min with PBS and incubated for at least 2 h in appropriate fluorophore-labelled secondary antibodies (donkey anti-rabbit AF488 (1:500, Invitrogen, A21206); donkey anti-guinea pig CF633 (1:500, Merck, SAB4600129)) diluted in blocking buffer. Sections were then washed 3 x 5 min with PBS before nuclei were stained with DAPI (1:5000, ThermoFisher, 62248) for 5 min, washed in PBS and mounted onto glass slides with Fluorescence Mounting Medium (Dako), and stored at 4◦C in the dark until imaging.

To stain meninges, dura mater was removed from brain into ice-cold PBS, fixed for 1 h in 4% buffered formaldehyde, and permeabilised in PBS containing 20mM EDTA for 30 min at 37 °C. Dura mater was blocked (3% BSA, 0.3% Triton-X100, 0.04% sodium azide in PBS) for 30 min at RT, before incubation in primary antibodies rabbit anti-IBA1 (1:500, Novachem, 019-19741) and rat anti-CD169 (1:200, BioLegend, 142402) diluted in blocking buffer overnight. Following this, dura mater was washed 3 x 5 min with PBS before incubation in secondary antibodies goat anti-rabbit AF594 (1:500, Invitrogen, A11012) and goat anti-rat AF647 (1:500, Invitrogen, A48265) diluted in blocking buffer for 1 h. Stained tissue was then mounted onto glass slides with Fluorescence Mounting Medium and stored at 4◦C in the dark until imaging.

Pregnant dams at E11.5 of both genotypes were culled and processed, and placenta and embryos were dissected. Embryo and placenta samples were immersion fixed in 4% buffered formaldehyde with 2% Triton-X100 for 30 min and washed 3 x 20 min in PBS. Tissues were then prepared for either cryosectioning (incubated in 30% sucrose in PBS for 24 h before they were embedded in OCT) or whole mount imaging. For whole mount imaging, whole embryos were cleared in RapiClear 1.52 (SUNJin Lab, RC152001) for 48 h at RT, before they were mounted in fresh RapiClear between two glass coverslips.

For cryosections, embryos with placenta were embedded in OCT, frozen and sectioned at 10 μm using a ThermoFisher HM525NX cryostat. Sections were rehydrated in Tris-buffered saline (TBS; 10mM Tris, 150mM NaCl, pH8.0) for 10 min and washed 3 x min in TBST (0.01% Tween-20 in

TBS). Blocking buffer (3% BSA, 5% serum in TBST) was applied for 30 min at RT, before sections were incubated with primary antibodies rabbit anti-RFP (1:100, Abcam, ab124754) and rat anti-TROMA-I (1:100, DSHB, TROMA-I) diluted in TBST for 1.5 h at RT. Sections were washed 3 x 5 min in TBST before incubation in secondary antibodies donkey anti-rabbit AF647 (1:500, Invitrogen, A31573) and donkey anti-rat AF488 (1:500, Invitrogen, A48269) diluted in TBST for 45 min protected from light. Sections were washed 3 x 5 min in PBS, stained with DAPI and mounted as above.

**Histochemistry**

Tissues (Liver, pancreas, spleen, small intestine) were fixed in 4% neutral buffered formalin and embedded in paraffin blocks. Sections were cut on a microtome (Leica) at 10µm, laid on a 40 ºC water bath and collected on Superfrost Plus slides (Bio-strategy). Slides were incubated at 60 ºC for 30 min before being de-waxed in xylene 2 x 5 min then rehydrated for 2 x 2 min washes in 100%, 95% and 75% ethanol, and washed in RO water for 5 min. Slides were stained with haematoxylin (Gill No 2, Sigma) for 30 seconds, rinsed under running tap water for 20 minutes and counterstained with Eosin Y (Sigma) for 40 seconds, dehydrated by 2 x 2 min washes in 70%, 95% and 100% ethanol, cleared 2 x 5 min in xylene and mounted with DPX mounting medium (Sigma).

For immunohistochemistry, antigen retrieval was performed on dewaxed and rehydrated sections, before incubation for 30 mins in 10 mM Tris/EDTA (pH 9) buffer maintained at 90-100ºC using a microwave. Slides were cooled on ice for 10 min, incubated in 0.3% hydrogen peroxide (Sigma) for 15 min to block endogenous peroxidases and washed in TBS. Slides were incubated in 1% BSA in TBS for 30 min followed by incubation with primary antibody (anti-F4/80, 1:1000, rabbit, Abcam) for 1.5 hr in a humidified chamber. Slides were washed with TBS and HRP secondary reagent (anti-rabbit, Agilent) applied for 30 min. Slides were washed again with TBS and then incubated for 5 min with DAB Peroxidase substrate (Agilent). Slides were washed with TBS and counterstained with haematoxylin (Gil No 2, Sigma) for 40 sec, then dehydrated and mounted as above. Imaging was performed on an Olympus BX50 microscope, using brightfield settings.

**Bone MicroCT**

MicroCT was performed on formalin-fixed samples and scanned using a Skyscan 1272 desktop MicroCT (Bruker, Belgium). A 7um voxel size was achieved using 4x4 camera binning and the following parameters: 70kV voltage, 142uA current, 1871ms exposure time, averaging of 2, Al 0.5mm filter, 0.5 degree rotation step and 360 degree rotation. The data was reconstructed with an FDK algorithm using NRecon 2.2.0.6 (Bruker, Belgium), and visualised with CTVox 3.3.1 (Bruker, Belgium). Volumetric analysis was performed using CTAn 1.20.8.0 (Bruker, Belgium). The trabecular region of interest was 2450um distal to the tibial metaphysis reference point and was 1750um in length.

**Quantification and assessment of the surface of osteoclasts**

Training images were created from the $TRAP^+$ data sets and loaded into a QuPath project to train the pixel classifier. $TRAP^+$, negative and background pixels were identified and the classifier trained until identifying $TRAP^+$ cells only. The classifier was applied against the whole well captures of the TRAP assays and used to segment the osteoclasts from the background and negative cells. QuPath was used to annotate the osteoclast TRAP+ cells with a minimum cut off above the size of the single cell population.

1. Keshvari S, Masson JJR, Ferrari-Cestari M, et al. Reversible expansion of tissue macrophages in response to macrophage colony-stimulating factor (CSF1) transforms systemic lipid and carbohydrate metabolism. *Am J Physiol Endocrinol Metab*. 2024;326(2):E149-E165.

#### Supplementary Table1

| Flow antibody list | | | |
|---|---|---|---|
| **Antibody** | **Flurorochrome** | **Reference** | **provider** |
| Anti-mouse CD11b | Percp-Cy5.5 | Cat#101228 | Biolegend |
| Anti-mouse Ly6G | BV785 | Cat#127645 | Biolegend |
| Anti-mouse Ly6C | BV421 | Cat#128008 | Biolegend |
| Anti mouse CD135 | APC | Cat#135310 | Biolegend |
| Anti mouse F4/80 | BUV395 | Cat#565614 | BD Biosciences |
| Anti mouse F4/80 | A647 | Cat#123122 | Biolegend |
| Anti mouse CD115 | BUV605 | Cat#135517 | Biolegend |
| Anti mouse c-kit | PECy7 | Cat#135112 | Biolegend |
| Anti-mouse CD11c | APC-Cy7 | Cat# 117323 | Biolegend |
| Anti-Mouse CD319 | BV711 | Cat#747994 | BD Biosciences |
| Anti mouse CD177 | AF647 | Cat#566599 | BD Biosciences |
| Anti mouse Ly-6A/E | BV650 | Cat#108143 | Biolegend |
| Anti mouse CD150 | BV711 | Cat#115941 | Biolegend |
| Anti mouse CD48 | BV421 | Cat#1103418 | Biolegend |
| Anti mouse CD16/32 | APC | Cat#101326 | Biolegend |
| Anti mouse CD34 | BV421 | Cat#343610 | Biolegend |
| Anti mouse CD71 | PE-Cy7 | Cat# 113812 | Biolegend |
| Anti mouse TER119 | APC | Cat#116212 | Biolegend |
| Biotin Anti-mouse Ly-6G/Ly-6C | | Cat# 108404 | Biolegend |
| Biotin Anti-mouse/human CD45R | | Cat# 103204 | Biolegend |
| Biotin Anti-mouse TER-119 | | Cat# 116204 | Biolegend |
| Biotin Anti-mouse CD3 Antibody | | Cat# 100244 | Biolegend |
| Biotin Anti-mouse CD5 Antibody | | Cat# 100604 | Biolegend |
| Streptavidin | APC-Cy7 | Cat# 405208 | Biolegend |

### Supp-Figure 1

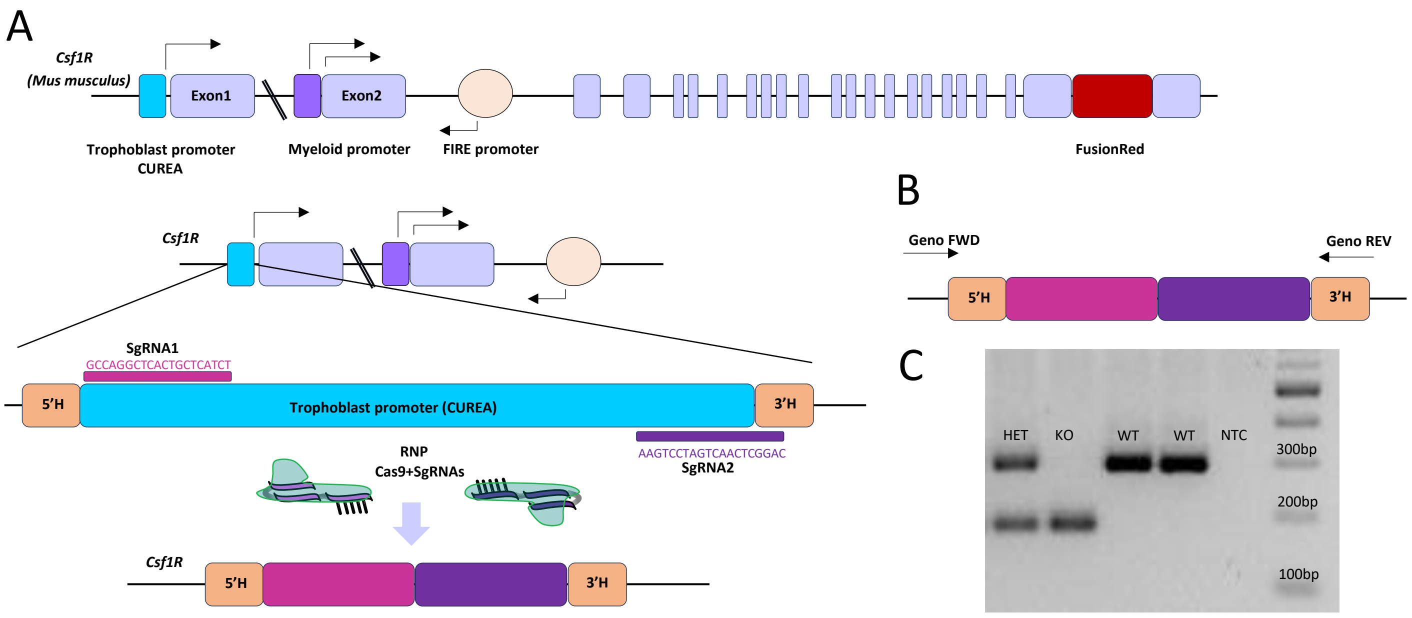

### Supp-Figure 2

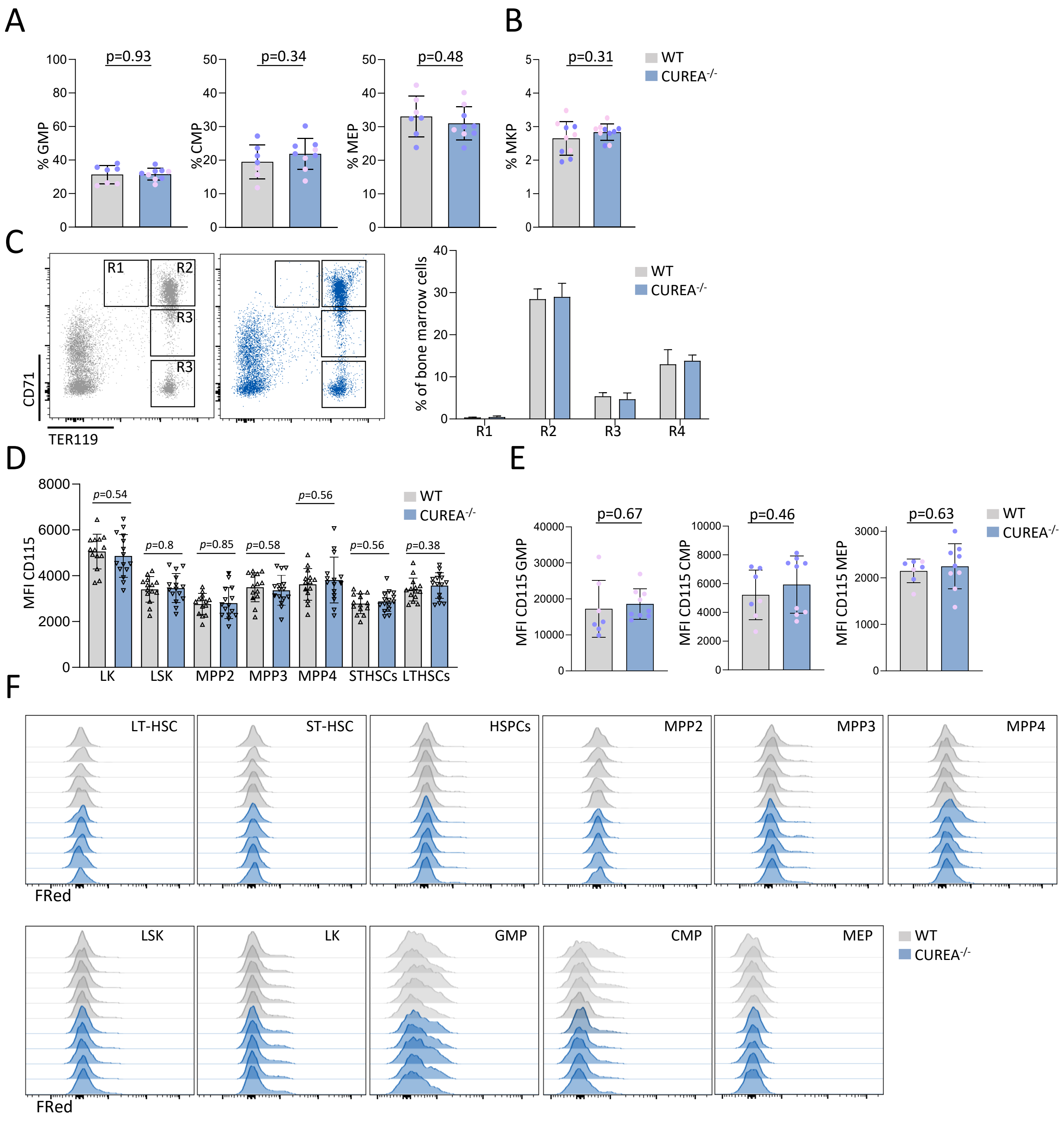

### Supp-Figure 3

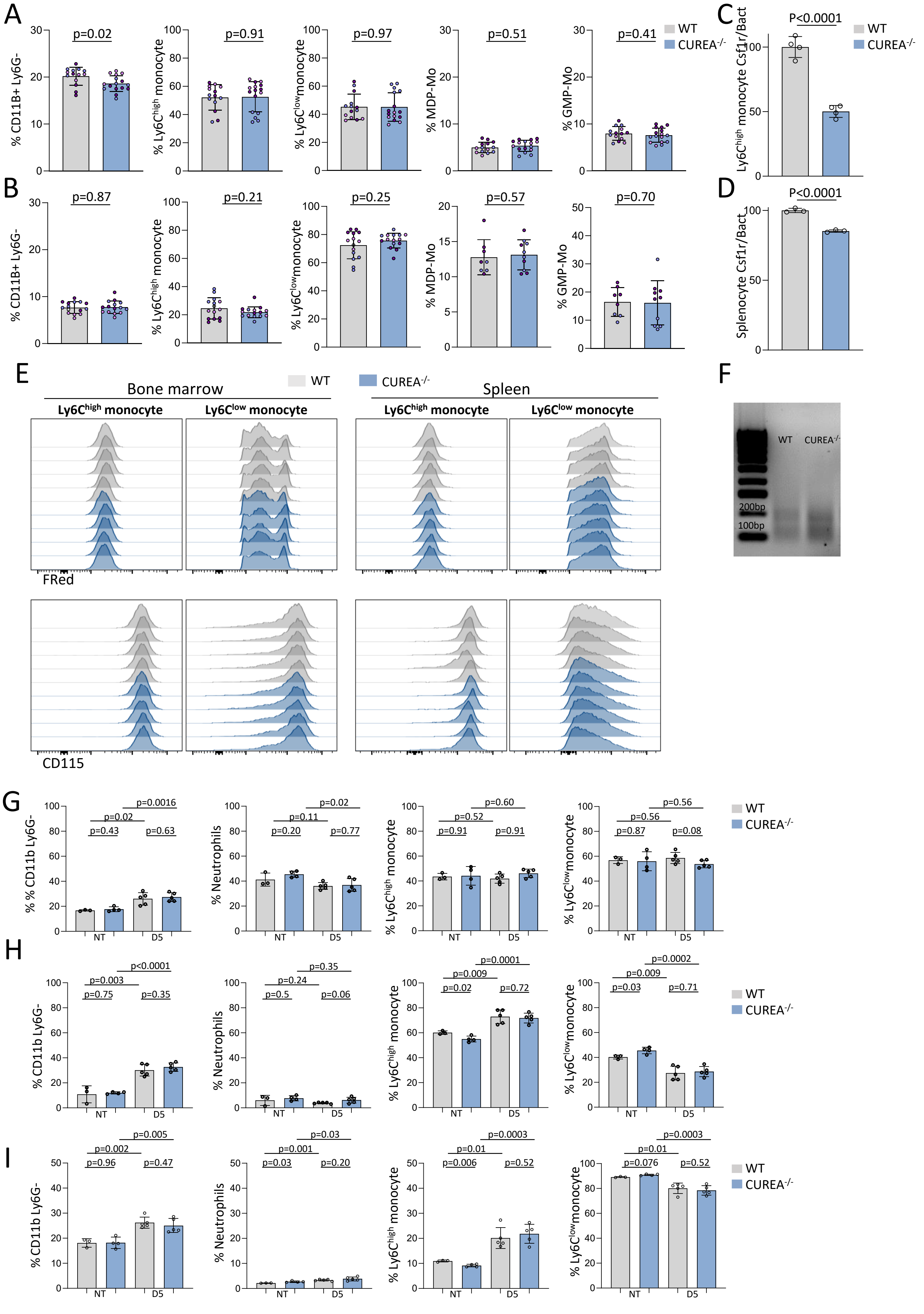

**Supp-Figure 4**

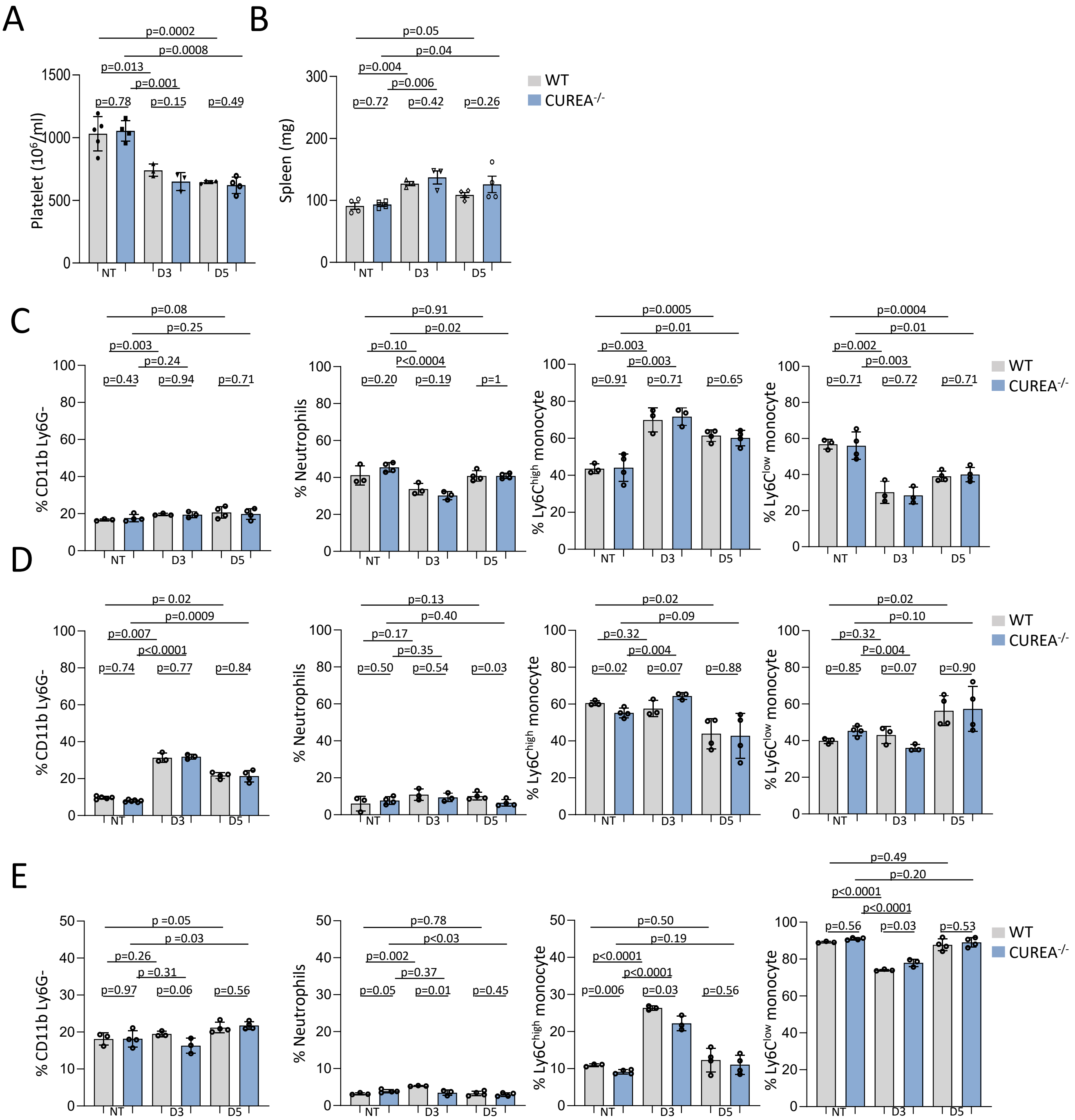

**Supp-Figure 5**

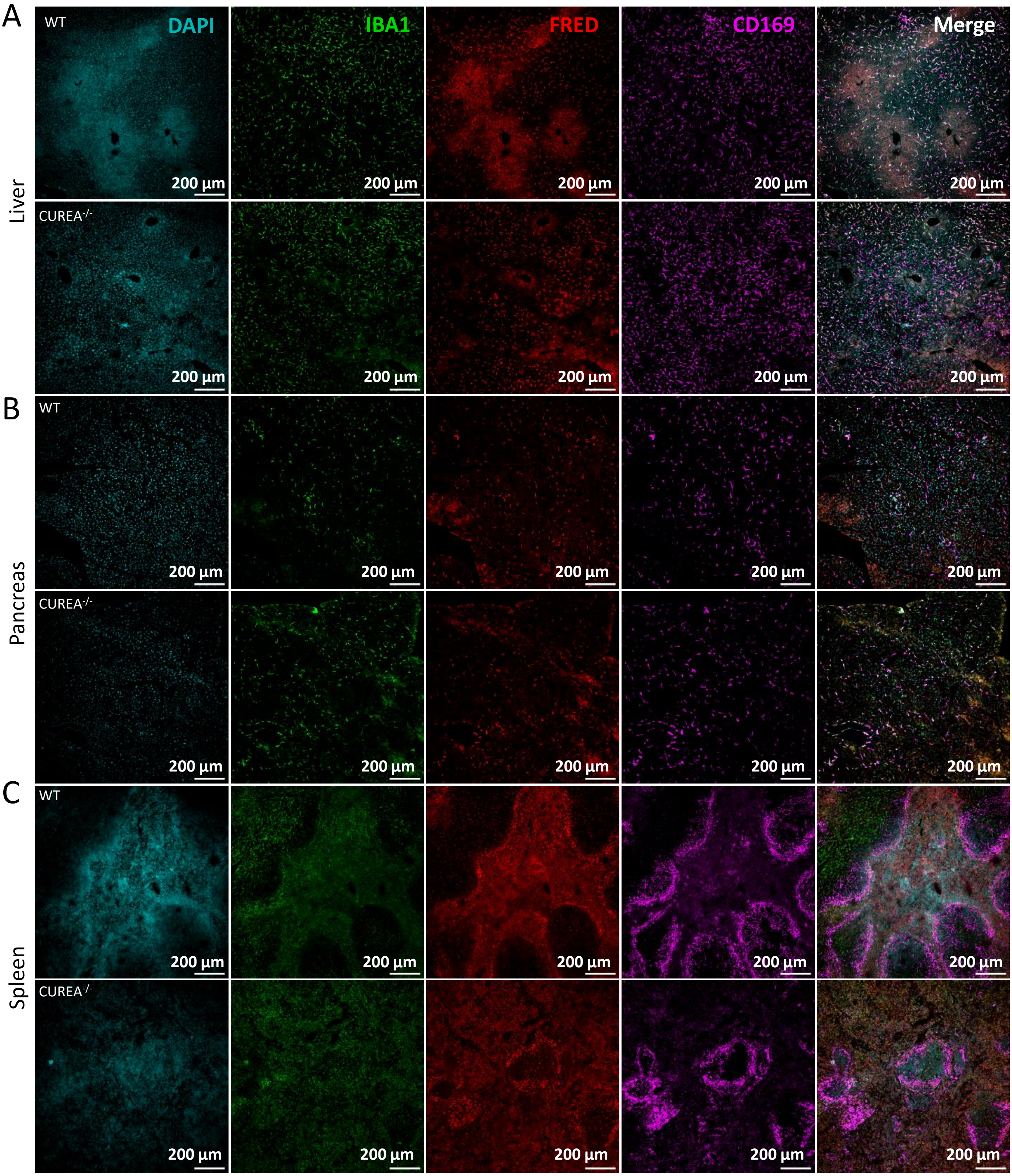

**Supp-Figure 6**

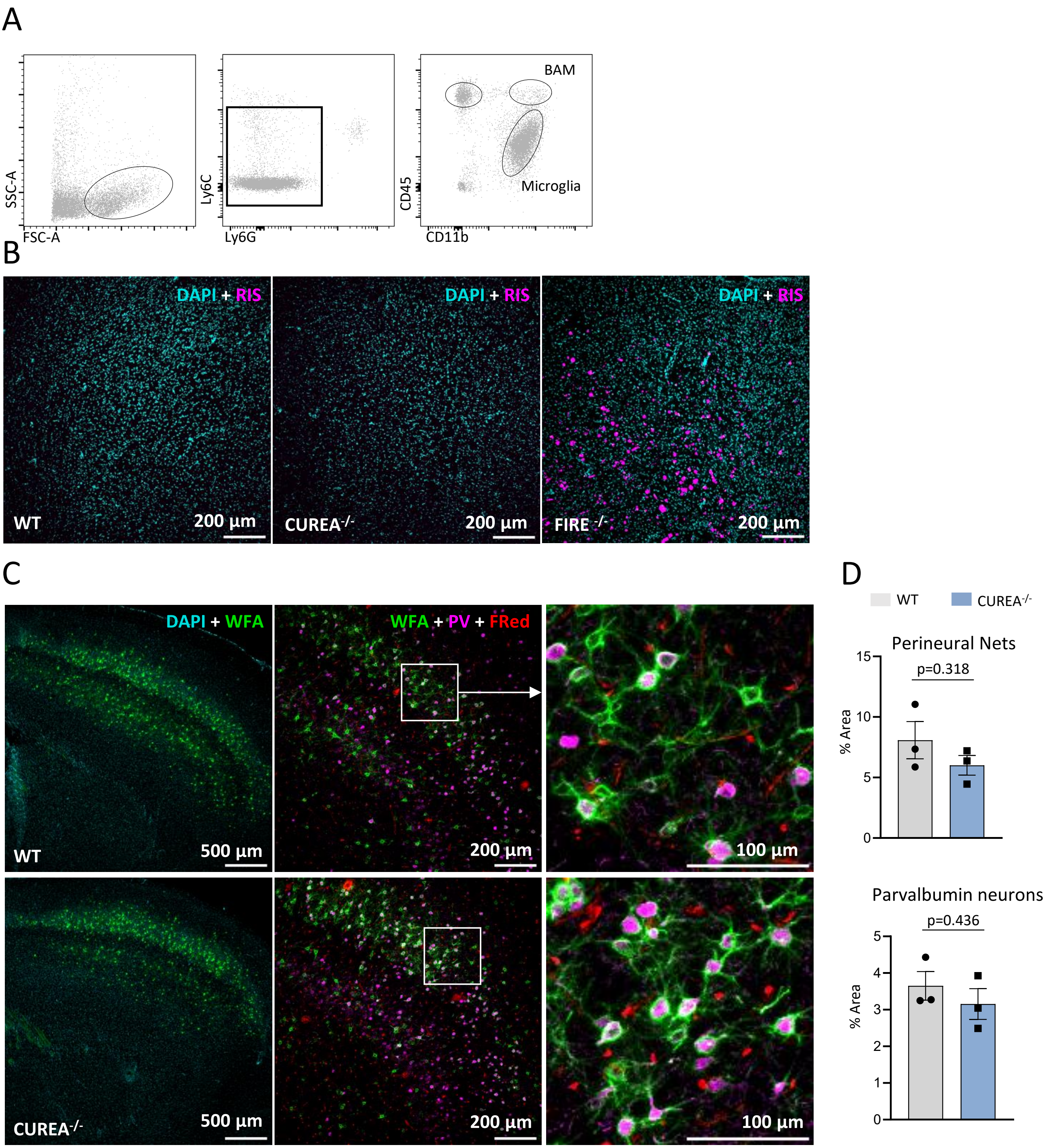

**Supp-Figure 7**

TGAGGAAACAGTTTGGTCCCCATGGCTCATATTTAGGGGTAACTTCATTTCTTCTCATGGGGTGA
CCGCTCTCACCTAGGGGGATAATGTCATCATGGAGTGATTTGTCTACAGGACTTGTTATTTTTGG
AACCTAGAATAAGTTTGGTTTAGGACAATTAGTGGAGACTTCCAGTTGCTGCTATAGGAGGCTCA
TTACTGTCAAAGTCCAAATCTCCAGTTACCTTGAGGCCTTCTGCCTTAAGGGCAGATGAGAAAGG
TATGAAGAATGTAGGATAACCTGACCTCTCTCAGCCAGGCTCACTGCTCATCTGGGAGACTC*TTA*
*TTTAT*GCTCCTGACTCAGAGGCTGCTGCTCTTGGGTATGACTCCTCTCCTAAGTGTCCTTAGAGG
TGGTTTCCTAAGTCTCTCAAACTCCATCATCTCCCTTCAGGATCAGTTGAGCCTGGCCCCAGATT
CTGCCTCTTCCTCTGGTGTGTGGTGTGTGTGTGTGTGTGTGTGTGTGTGTGTGTGTATGTGTG
TGTGCTTTGTTTTCTTCTAGAGACCCAATATTTCCAAATTCTGTAGTTCCCTTTCAGGCAACCTA
AAAAAAAAAAAAAAAAAAAAGGGGAAGAGGAGCCAGTGCAACAGACAGGAACGTGTTCATCTGTT
CCCGTCCTCACAGAACTAGCAGCTGGGAGCCCCGTGCCCAGCCGACTCTCCAACCTGCATCGGCT
CACGCTATCCCCTGGAGGCTATGGAGTTGGGGCCTCCTCTGGTCCTGCTGCTGGCCACAGTTTGG
CATG

CUREA *TATA box* F1.1. F1.2 R2. Myeloid TSS Trophoblast TSS

splice acceptors. splice donors.
